# Population sequencing reveals clonal diversity and ancestral inbreeding in the grapevine cultivar Chardonnay

**DOI:** 10.1101/389197

**Authors:** Michael J. Roach, Daniel L. Johnson, Joerg Bohlmann, Hennie J.J. van Vuuren, Steven J. M. Jones, Isak S. Pretorius, Simon A. Schmidt, Anthony R. Borneman

## Abstract

Chardonnay is the basis of some of the world’s most iconic wines and its success is underpinned by a historic program of clonal selection. There are numerous clones of Chardonnay available that exhibit differences in key viticultural and oenological traits that have arisen from the accumulation of somatic mutations during centuries of asexual propagation. However, the genetic variation that underlies these differences remains largely unknown. To address this knowledge gap, a high-quality, diploid-phased Chardonnay genome assembly was produced from single-molecule real time sequencing, and combined with re-sequencing data from 15 different commercial Chardonnay clones. There were 1620 markers identified that distinguish the 15 Chardonnay clones. These markers were reliably used for clonal identification of validation genomic material, as well as in identifying a potential genetic basis for some clonal phenotypic differences. The predicted parentage of the Chardonnay haplomes was elucidated by mapping sequence data from the predicted parents of Chardonnay (Gouais blanc and Pinot noir) against the Chardonnay reference genome. This enabled the detection of instances of heterosis, with differentially-expanded gene families being inherited from the parents of Chardonnay. Most surprisingly however, the patterns of nucleotide variation present in the Chardonnay genome indicate that Pinot noir and Gouais blanc share an extremely high degree of kinship that has resulted in the Chardonnay genome displaying characteristics that are indicative of inbreeding.

**Author Summary:** Phenotypic variation within a grapevine cultivar arises from an accumulation of mutations from serial vegetative propagation. Old cultivars such as Chardonnay have been propagated for centuries resulting in hundreds of available ‘clones’ containing unique genetic mutations and a range of various phenotypic peculiarities. The genetic mutations can be leveraged as genetic markers and are useful in identifying specific clones for authenticity testing, or as breeding markers for new clonal selections where particular mutations are known to confer a phenotypic trait. We produced a high-quality genome assembly for Chardonnay, and using re-sequencing data for 15 popular clones, were able to identify a large selection of markers that are unique to at least one clone. We identified mutations that may confer phenotypic effects, and were able to identify clones from material independently sourced from nurseries and vineyards. The marker detection framework we describe for authenticity testing would be applicable to other grapevine cultivars or even other agriculturally important woody-plant crops that are vegetatively propagated such as fruit orchards. Finally, we show that the Chardonnay genome contains extensive evidence for parental inbreeding, such that its parents, Gouais blanc and Pinot noir, may even represent first-degree relatives.

## Introduction

Chardonnay is known for the production of some of the world’s most iconic wines and is predicted to be the result of a cross between the *Vitis vinifera* cultivars Pinot noir and Gouais blanc (1, 2). Since first appearing in European vineyards, Chardonnay has spread throughout the world and has become one of the most widely cultivated wine-grape varieties (3). For much of the 20^th^ century, grapevine cultivars were generally propagated by mass selection. High genetic variability therefore existed between individual plants within a single vineyard and this heterogeneity often lead to inconsistent fruit quality, production levels, and in some wine-producing regions, poor vine health (4). Clonal selection arose as a technique to combat these shortcomings, preserving the genetic profile of superior plants, while amplifying favourable characteristics and purging viral contamination, leading to improved yields (4, 5).

Chardonnay’s global expansion throughout commercial vineyards, which started to accelerate rapidly in the mid-1980s, coincided with the maturation of several clonal selection programmes based in France, the USA and Australia. As a result, there are now many defined clones of Chardonnay available that exhibit differences in key viticultural and oenological traits (3, 6–11). For example, clone I10V1—also known as FPS06 (12)—showed early promise as a high-yielding clone with moderate cluster weight and vigorous canopy (13). The availability of virus-free clonal material of I10V1 helped cement productivity gains in the viticultural sector and I10V1 quickly dominated the majority of the Australian Chardonnay plantings (4, 5).

Since the concurrent publication of two draft Pinot noir genomes in 2007 (14, 15) grapevine genomics has increasingly contributed to the understanding of this woody plant species. However, the haploid Pinot noir reference genome does not fully represent the typical complexity of commercial wine-grape cultivars and the heterozygous Pinot noir sequence remains highly fragmented (16). In recent years, the maturation of single molecule long-read sequencing technology such as those developed by PacBio (17) and Oxford Nanopore (18), and the development of diploid-aware assemblers such as FALCON (19) and CANU (20) has given rise to many highly-contiguous genome assemblies, including a draft genome assembly for the grapevine variety Cabernet sauvignon (19, 21–24). Furthermore, whole genome phasing at the assembly level is possible with assemblers such as FALCON Unzip (19), allowing both haplotypes of a diploid organism to be characterised. For heterozygous diploid organisms, such as Chardonnay, this is especially important for resolving haplotype-specific features that might otherwise be lost in a traditional genome assembly.

The aim of this work was to explore the diversity extant within Chardonnay clones. A reference genome for Chardonnay was assembled *de novo* from PacBio long-read sequence data against which short-read clonal sequence data was mapped. This led to the identification of clone diagnostic single-nucleotide polymorphisms (SNP) and Insertions/Deletions (InDel) that show little shared clonal heritage. Furthermore, comparison of the Chardonnay reference with Pinot noir revealed some unexpected complexities in haplotype features with implications for the pedigree of this important grapevine variety.

## Results

### Assembly and annotation of a high quality, heterozygous phased Chardonnay genome

Of the many Chardonnay clones available, clone I10V1 was chosen as the basis for the reference genome due to its prominent use in the Australian wine industry. The initial I10V1 genome was assembled, phased and polished using subreads generated from 54 PacBio RS-II SMRT cells and the FALCON Unzip, Quiver pipeline (19, 25). While this assembly method should produce an assembly in which the primary contigs represent the haploid genome content of the organism in question, the size of the initial assembly (580 Mb) significantly exceeded that expected for *V. vinifera* (450–500 Mb). Both analysis with BUSCO (26) and short-read mapping indicated that this increased size was primarily due to both copies of many genomic regions (rather than only a single haplotype) being represented in the primary contigs (S1 Table and S2 Fig), a situation that is common in heterozygous diploid genome assemblies (19, 27–29). To address these assembly artefacts, the initial primary contig pool was aggressively de-duplicated, with small primary contigs that were allelic to larger primary contigs being re-assigned to the haplotig pool. This approach reassigned 694 primary contigs (100 Mb) and added 36 haplotigs (11 Mb), while also purging 18 repeat-rich artefactual contigs (1.3 Mb). Manual curation, based upon alignments to the PN40024 assembly (14) and subread mapping were used to address several remaining mis-assemblies.

The final curated Chardonnay assembly consists of 854 primary contigs (N_50_ of 935 kb) and 1883 haplotigs, totalling 490 Mb and 378 Mb, respectively (Table 1). There were approximately 95% complete, and only 1.6% fragmented BUSCO-predicted genes (Supplementary Table S1). BUSCO duplication is also predicted to be low for both the primary contigs and the associated haplotigs (4% and 2% respectively). A custom repeat library was constructed for Chardonnay and used to annotate 336 Mb (38.7%) of the diploid genome as repetitive. RNAseq data were used to annotate potential coding regions of the primary contigs using Maker (30), which predicted 29 675 gene models (exclusive of repetitive regions) and 66 548 transcripts in total.

**Table 1:** Quast-based assembly statistics for the Chardonnay clone I10V1 genome

|  | Primary contigs | Haplotigs |
| --- | --- | --- |
| Contigs | 854 | 1883 |
| Contigs ($\geq 50$ kb) | 838 | 1614 |
| Assembly Size (Mb) | 490.0 | 378.0 |
| Largest Contig (Mb) | 6.35 | 1.91 |
| GC (%) | 34.41 | 34.45 |
| N50 (kb) | 935.8 | 318.4 |
| N75 (kb) | 502.7 | 165.3 |
| L50 | 145 | 335 |
| L75 | 321 | 749 |

### Phasing coverage, and identification of homozygous and hemizygous regions

A total of 614 primary contigs (397 Mb) and 1502 haplotigs (305 Mb) were confidently placed in chromosomal order using the PN40024 scaffold as a reference. To analyse the degree and distribution of heterozygosity across the genome, read depth (from mapped RS II subreads), heterozygous variant density (from mapped Illumina short-reads) and phasing coverage (from contig alignments) was calculated for the assembly (Fig 1A).

**Fig 1.**
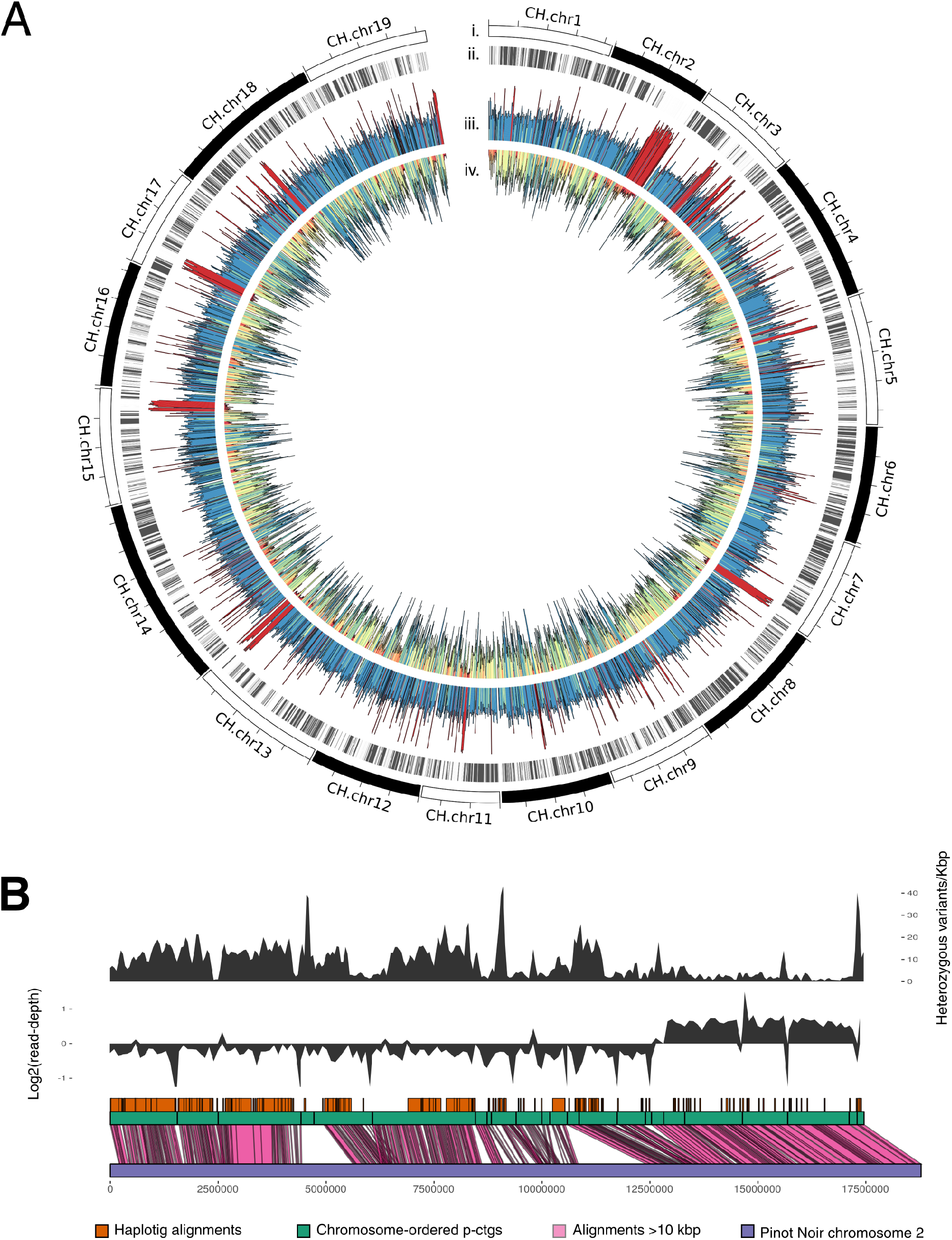
The *Vitis vinifera* cultivar Chardonnay reference genome. (*A*) A circos plot showing chromosome-ordered primary contigs (*i*), haplotig alignments (*ii*), read-depth of RS II subreads mapped to diploid assembly (read-depth colour scale: yellow, low; blue, high; red, double) (*iii*), and heterozygous variant density (SNP density colour scale: red, low; blue, high) (*iv*). (*B*) An expanded view of Chardonnay Chromosome 2 showing heterozygous variant density (*top* track), log2 read-depth (*middle* track), and alignment with Pinot noir Chromosome 2 (*bottom* track).

Chromosomes 2, 3, 7, 15, 17, and 18 contain runs of homozygosity greater than 500 kb (intersect of lack of phasing coverage, double read-depth, low heterozygous variant density). There are a further 22.8 Mb that lacked phasing coverage, had low heterozygous variant density, and median read-depth. These regions presumably result from either hemizygosity of these genomic regions, or undetected allelic duplicates remaining in the primary contigs.

The largest homozygous run identified resides on Chromosome 2 and aligns closely to the Pinot noir assembly at over 99.8% identity (Fig 1B). A region of synteny remaining in the primary contigs is present (CH.chr2:5570000– 6520000), evidenced by the ends of neighbouring primary contigs aligning to the same region in Pinot noir. In addition, there were two regions of low heterozygous variant density, poor phasing coverage, and median read-depth (CH.chr2:99000000–10300000 and CH.chr2:11450000–12600000). BLAST searches for these regions within the remaining primary contigs and haplotigs did not reveal any significant alignments. As such these regions appear to be hemizygous.

### Defining parental contributions to the Chardonnay genome

To further refine the relationship between Chardonnay and the two varieties previously reported to be its parents (1, 2, 31), an attempt was made to identify the parental origin of each allele in the diploid Chardonnay assembly. Phase blocks were assigned across the genome by aligning and trimming both the primary contigs and haplotigs into pairs of syntenic sequence blocks (P and H alleles). This produced 1153 phase-blocks covering 270 Mb of the genome (55%). Each pair of phase blocks should have one allele inherited from each parent. To assign likely genomic parentage within each phase block, short-reads from Gouais blanc, and a merged dataset comprising sequencing reads from several different Pinot varieties (32) (hereafter referred to as Pinot) were mapped to the phase block sequences. The proportion of inherited nucleotide variation (using heterozygous variant loci) was then used to attribute the likely parentage of each block.

It was possible to confidently assign parentage to 197 Mb of the 244 Mb of chromosome-ordered phase-blocks (Fig 2A). Interestingly, rather than a 1:1 ratio of Gouais blanc to Pinot matches, Pinot was shown to match a higher proportion of the phase blocks (49% versus 34% Gouais blanc), suggesting that the Pinot noir genome has contributed a higher proportion of genetic material to Chardonnay than Gouais blanc. However, further complicating this imbalance was the observation that in the remaining 17% of assigned regions, the pattern of nucleotide variation across the two heterozygous Chardonnay haplotypes matched both haplotypes of Pinot, with one of these haplotypes also matching one of the Gouais haplotypes. These ‘double Pinot haplotype’ regions are in some cases many megabases in size and are indicative of a common ancestry between Pinot and Gouais blanc.

**Fig 2.**
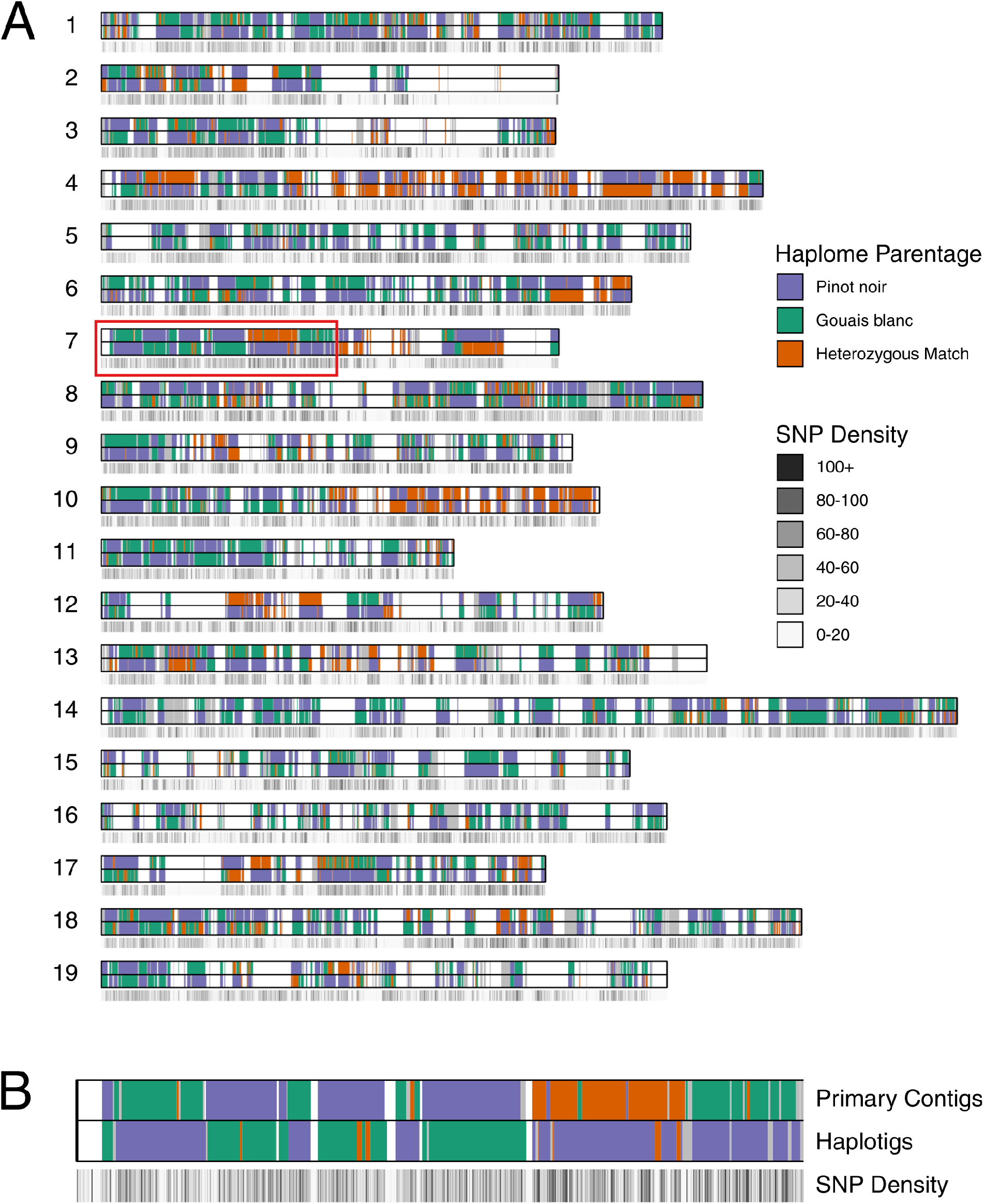
Parental architecture of the Chardonnay genome. (*A*) An ideogram of the Chardonnay reference assembly with the positions of both primary contig and haplotig phase-blocks indicated and juxtaposed with a SNP density track (for the primary contigs). Gaps in phase-blocks are indicated in white. (*B*) An enlargement of a region of *Vitis vinifera* Chromosome 7 (red box) in Figure 2A.

While reciprocity (one Gouais blanc haplotype, one Pinot haplotype) was observed between allelic phase-blocks for over 95% of the parentage-assigned sequence, frequent haplotype switching (a known characteristic of FALCON-based assemblies) was observed between the haplomes, producing a haplotype mosaic which is observable as a ‘checkerboard’ pattern that alternates between the primary contigs and haplotigs for each chromosome (Fig 2B).

### Parental-specific genomic variation

With parental contributions delineated in the Chardonnay assembly, it was possible to determine the parental origins of structural variation between orthologous chromosomes, including parent-specific gene family expansions. Tandem pairs of orthologous proteins were defined in Chardonnay and filtered to identify tandem orthologs that were both expanded in Chardonnay compared to the Pinot noir reference assembly, and which resided in the Gouais blanc haplome. Chromosome alignments containing gene expansion candidates were inspected for features consistent with tandem gene duplication. Using this analysis, an expansion of Fatty Acyl-CoA Reductase 2-like (*FAR2-like*) genes on Chromosome 5 was identified, with the arrangement of *FAR2-like* open reading frames (ORFs) consistent with a tandem duplication event (S3 Fig).

A protein-based phylogeny was produced that encompassed the four *FAR2-like* ORFs present in the Chardonnay assembly, in addition to the homologous proteins from the Pinot noir PN40024 assembly (Fig 3A). Using these data, the Chardonnay haplotig sequence was identified as being derived from Pinot (nucleotide sequences of *FAR2-like* genes from Pinot noir and the Chardonnay haplotig were identical). However, rather than having an orthologous set of protein-coding regions, the genomic sequence derived from Gouais blanc (present in the primary contig) is predicted to encode two additional copies of *FAR2-like* homologues and an extra *FAR2-like* pseudogene (Fig 3B). While the ORF that was orthologous to the Pinot *FAR2-like* gene was closely related to the Pinot noir *FAR2-like* gene (98% identity), the two additional ORFs from Gouais blanc were more distantly related (93– 94% identity), suggesting that this gene expansion was not a recent event.

**Fig 3.**
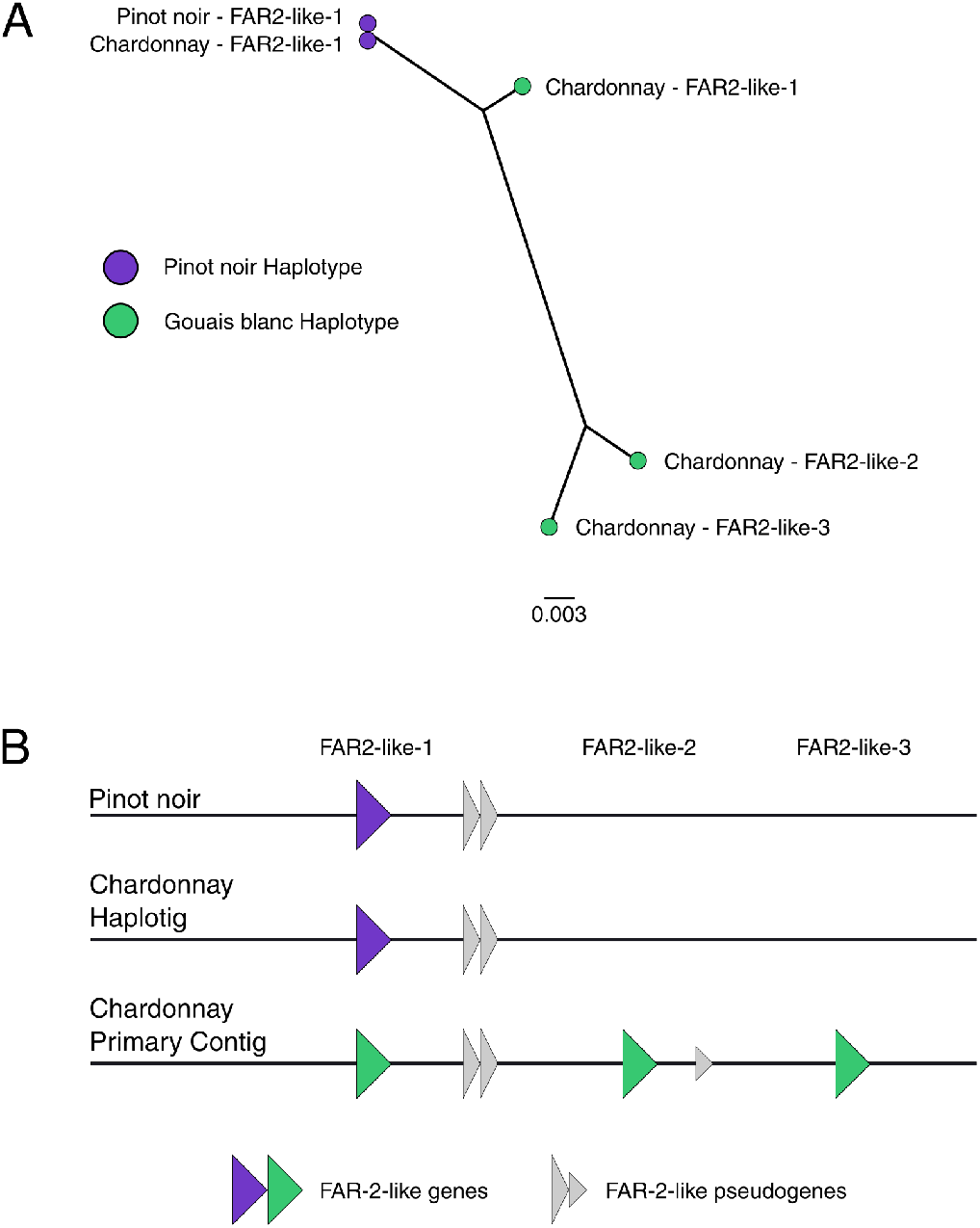
A FAR2-like expanded gene family in Chardonnay. (*A*) An unrooted tree of *FAR2-like* genes. (*B*) A schematic of the predicted genomic arrangement of the *FAR2-like* genes in Chardonnay. Both Pinot noir-derived (purple) and Gouais blanc-derived genes (green) are shown.

### Clonal nucleotide variation within a grapevine cultivar

As for many commercial grapevine varieties, there are currently many clones of Chardonnay, with each exhibiting a unique range of phenotypic traits. However, unlike varietal development, all of these genetic clones were established through the repeated asexual propagation of cuttings that presumably trace back to an original Chardonnay plant. It is therefore an accumulation of somatic mutations, that has contributed to phenotypic differences that uniquely define each clone and which provide an avenue for the confirmation of a clone’s identity. While clonal variation has so far been ill-defined in grapevine, the availability of the Chardonnay reference genome provides an opportunity to investigate the SNP spectrum that has arisen during the long history of Chardonnay propagation.

To begin to catalogue the diversity that exists across the clonal landscape of Chardonnay, short-read re-sequencing was used to define single nucleotide variation across 15 different Chardonnay clones. The analysis of these highly related genomes (separated by a low number of true SNPs) was facilitated through the use of a marker discovery pipeline developed to call variants while applying a stringent kmer-based filter to remove false positives (including those calls due to sequencing batch or individual library size distribution at the expense of some false negative calls). Similar kmer approaches have been reported with excellent fidelity (33). After filtering, 1620 high confidence marker variants were identified and evenly distributed across the Chardonnay genome (Table 2, S4 Fig, and Sheet 1 in S5 Dataset). Variant calls were concatenated and used to generate a Chardonnay clone phylogeny (Fig 4).

**Table 2:** A summary of Chardonnay clonal marker variants

| Sample Group | Number of SNPs and InDels |
| --- | --- |
| Mendoza | 221 |
| 809 | 187 |
| 95 | 150 |
| G9V7 | 143 |
| 277 | 137 |
| 76 | 137 |
| 548 | 133 |
| 352 | 121 |
| 1066 | 120 |
| 96 | 61 |
| CR Red, Waite Star, I10V1 | 60 |
| 118 | 27 |
| Waite Star | 26 |
| 118, 352, 124 | 24 |
| CR Red, Waite Star, I10V1, 277 | 18 |
| CR Red | 14 |
| 352, G9V7 | 11 |
| 76, 548 | 10 |
| CR Red, Waite Star, I10V1, Mendoza, G9V7, 95, 277, 352, 96, 1066, 124, 118, 809 | 8 |
| 118, 124, 96 | 4 |
| CR Red, Waite Star, I10V1, Mendoza, 809, 95, 277 | 2 |
| CR Red, I10V1 | 2 |
| 118, 1066, 124, 96 | 2 |
| 124 | 2* |

\*alternate base calls were very low for these two variants.

**Fig 4.**
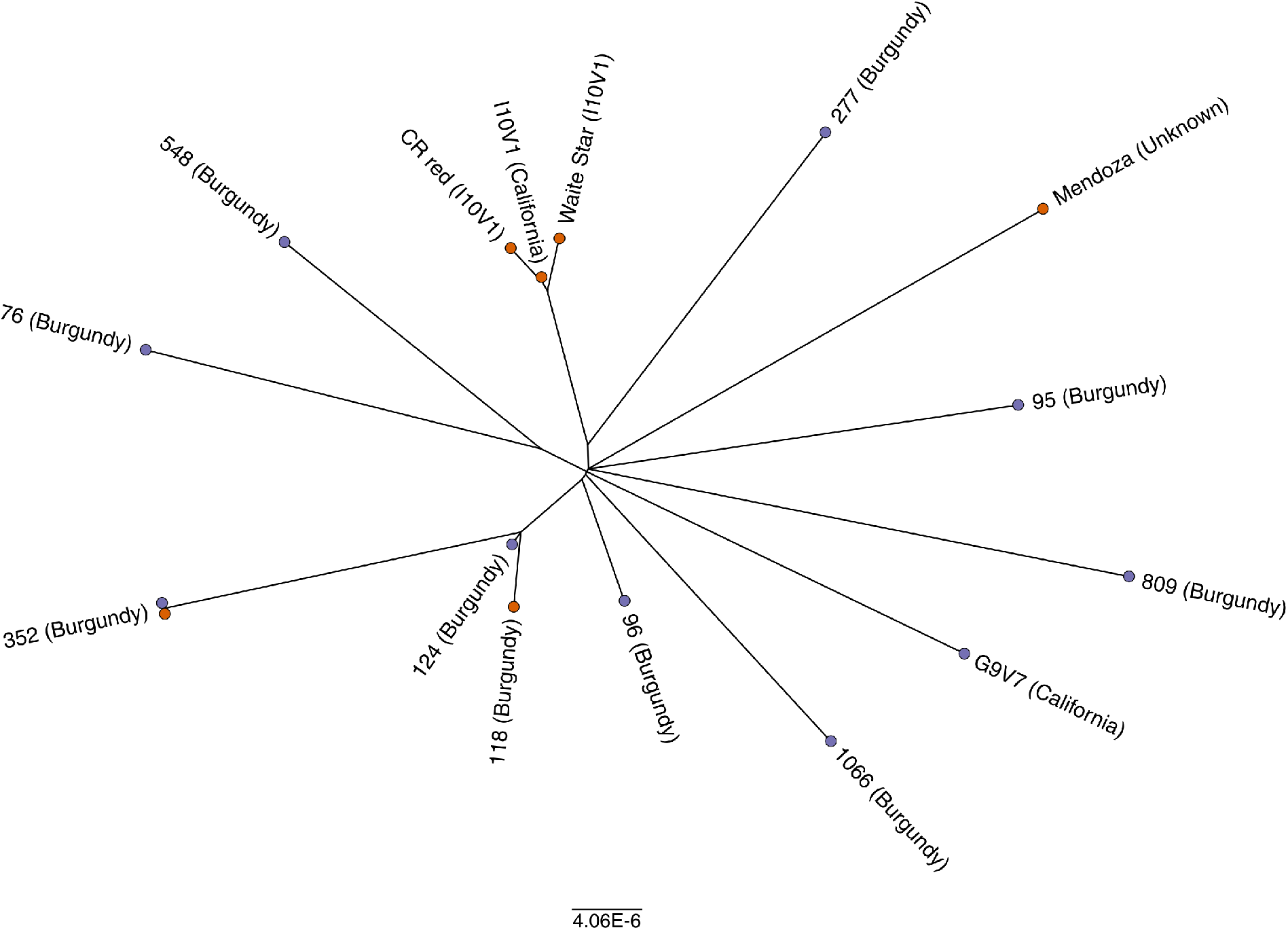
Genetic diversity in Chardonnay clones. An unrooted tree of Chardonnay clones based upon bi-allelic SNPs. Sequencing batches are designated by coloured terminal nodes (orange, sequencing batch *#*1; purple, sequencing batch *#*2).

‘CR Red’ and ‘Waite Star’, suspected phenotypic mutants of I10V1 that have red-skinned and seedless berries respectively (34, 35), formed a tight clade with only 40 variants (36 SNPs, 4 InDels) separating the three samples. The tight grouping of these clones confirms that the variant discovery pipeline can reliably detect recent clonal relationships from independent tissue samples. There were no further *a priori* relationships known for the remaining clones. However, the variant analysis would suggest that clones 124 and 118 also share some common ancestry as they are separated by only 23 SNPs.

The accumulation of SNPs can also lead to phenotypic differentiation that underlies the clonal selection process. For example, the major clonal-specific phenotypic variant of Chardonnay, “Muscat character”, results from one of several single nucleotide substitutions that produce non-synonymous amino acid changes in 1-deoxy-D-xylulose-5-phosphate synthase 1 (*DXS1*) gene and are associated with the production of higher levels of monoterpenoids (36). A combination of Annovar (37) and Provean (Choi *et al.,* 2012) were therefore used to annotate and predict the potential protein-coding consequences of each of the marker variant mutations identified. This pipeline correctly identified a previously characterised Muscat mutation (*S272P*) in *DXS1* in clone 809, the only Chardonnay clone in this study known to display the Muscat character. Provean scored this mutation at –3.37, where values less than −2.5 generally signify an increased likelihood that the mutation impacts the function of the enzyme. In addition to this known Muscat mutation, an additional 55 marker mutations were identified that displayed a high chance of impacted protein function (Sheet 2 in S5 Dataset). However, further work is required to investigate the links between known inter-clonal phenotypic variation and these specific mutations.

### The application of SNP and InDel-based markers for clone-specific genotyping

While various phenotypic characteristics (known as ampelography) and microsatellite based genetic tests can be used to positively identify grapevines, the accurate identification of specific clones is extremely difficult and to date, microsatellite-based marker systems have proven unreliable for the identification of clonal material (38, 39). Uncertainties can therefore exist as to the exact clone that has been planted in many vineyards. To enable a rapid clonal re-identification methodology, a kmer-approach was developed (similar to the method described in Shajii, Yorukoglu (40)) for screening raw short-read sequence data from unknown Chardonnay samples against the pre-identified clonal-specific variants. This method queries known marker variants against a kmer count database generated from the unknown sample. The matching markers and sample groups are returned allowing the potential identification of the unknown Chardonnay sample.

The marker detection pipeline was tested using data from a variety of different samples and sequencing methods (Sheet 3 in S5 Dataset). Chardonnay clones 76, 124, 548, 809, 1066, and G9V7 were independently sequenced at a second location from the same genomic DNA that was used for the original identification of nucleotide variants; however, both a different library preparation and sequencing platform were used. This enabled an evaluation of marker suitability and false discovery rates in a best-case scenario (i.e. when the source DNA was the same). When screened with the pipeline (Fig 5A), between 29% and 76% of the markers were detected for each of the samples and nearly all the missing markers coincided with poor coverage at marker loci.

**Fig 5:**
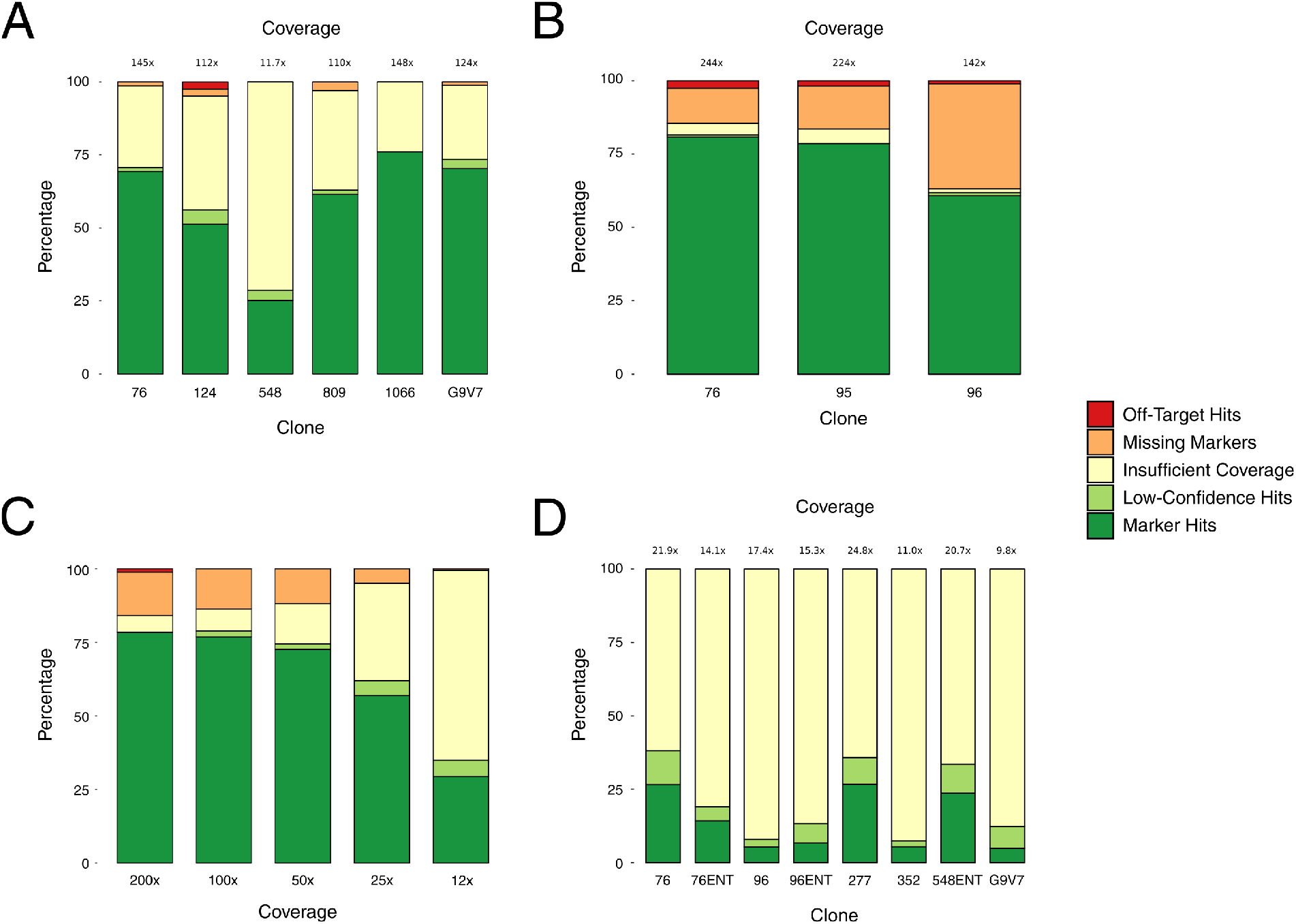
Marker screening using WGS data in Chardonnay. (*A*) Moderate coverage sequencing of clonal material used in marker discovery. (*B*) High-coverage sequencing of independently-sourced clonal material. (*C*) Subsampling of High-Coverage, independently-sourced clonal material (clone 95 from Figure 5B). (*D*) Low-coverage sequencing of independently-sourced clonal material (‘ENT’ denotes ENTAV-INRA^®^ source material).

To validate suitability of the markers for clonal identification, Chardonnay clones 76, 95, 96, 277, 352, 548 and G9V7 were independently sourced and sequenced. High-coverage (142- to 244-fold) sequencing was performed for three of these independently-sourced clones (Fig 5B). Kmer analysis identified between 62% and 81% of the expected markers for each sample, with minimal (1.2% to 2.6%) off-clone variants detected. However, despite being the same clones, there were a significant proportion (12% to 36%) of the expected markers in each of these three samples that were not found in the independent material and which could not be attributed to insufficient marker loci coverage, which indicates that there may be intra-clonal genetic variation that has accumulated during the independent passaging of clonal material.

As the level of sequencing coverage ultimately impacts the economics of clonal testing, the impact of sequencing depth on marker identification was assessed. Data consisting of the pooled results of two sequencing batches for independently-sourced clone 95 was subsampled to a range of coverages and then screened for clonal identification effectiveness (Fig 5C). At 200-fold coverage there were only 2 (low confidence) off-target hits, and none at lower coverages. There was little difference in the number of discoverable markers from 200-fold down to 25-fold coverage (79% and 73% respectively), and only a 4% decrease in markers confidently-flagged as missing. At 12-fold coverage it was still possible to detect 58% of the markers for this clone.

Given the successful results of the coverage titration, low coverage (9.8- to 24.8-fold) datasets were obtained from independent material of six clones, with clones 76 and 96 each sourced from proprietary and generic selections (Fig 5D). Despite the combination of independent material and low coverage it was still possible to detect between 7% and 38% of the expected markers for each sample, with no off-target hits.

## Discussion

The genomic complexity of grapevine, combined with its clonal mode of propagation (absence of outcrossed populations), has so far limited classical genetic approaches to understanding inherited traits in this valuable crop. The availability of a reference genome for *V. vinifera* (14, 15) has facilitated genetic studies through the provision of additional microsatellite markers for parentage and other studies (31, 41–43) and more recently has driven the development of dense SNP arrays that are being used for analysis of population structure and genome wide association studies (44–46). While not subject to the same technical limitations of microsatellite analysis (47), using predefined sets of SNPs also has its limitations, particularly with regard to discovery of novel genomic features. Recent advances in sequencing technology, and specifically read length, have provided a way forward, enabling repeat-rich genomes, such as grapevine, to be considered in their native state, without having to strip its inherent genomic variability in order to achieve a genome model with moderate contiguity.

A reference genome for Chardonnay was produced using long-read single-molecule sequence data in order to more precisely and accurately define the differences between the almost identical derivatives (clones) of a single cultivar. The Chardonnay assembly reported here exhibits a high level of contiguity and predicted completeness and provides a fundamental platform for the in-depth investigation of Chardonnay’s genome function and, more generally, of grapevine evolution and breeding.

Heterosis has been reported to have played a large role in the prominence of Gouais blanc and Pinot noir crosses in wine grapevines (5). Deleterious mutations in inbred lines can lead to increased susceptibility to pests and diseases, reduced stress tolerance, and poorer biomass production (5). This can be offset with the introduction of novel genes and gene families by crossing with a genetically dissimilar sample. The inheritance of an expanded family of *FAR2-like* genes from Gouais blanc represents one example of where this may have occurred in Chardonnay. The sequence divergence in *FAR2-like* copies and haplotypes suggests that the gene expansion event was not a recent occurrence. The increased gene copy number and sequence diversity potentially enriches the Chardonnay genome for both redundancy and functionality of this gene.

Fatty Acyl-CoA Reductase (FAR) enzymes catalyse the reaction: long-chain acyl-CoA + 2 NADPH → CoA + a long-chain alcohol + 2 NADP^+^ (48, 49). There are numerous copies of FARs in plants and each tends to be specific for long-chain acyl-CoA molecules of a certain length (50). FARs form the first step in wax biosynthesis and are associated with many plant surfaces, most notably epicuticular wax. Epicuticular waxes are important for protecting plants against physical damage, pathogens, and water loss (51–55). It was reported in Konlechner and Sauer (56) that Chardonnay has a very high production and unique pattern of epicuticular wax; this might be attributed to novel FARs. The fatty alcohols produced by *FAR2* are associated with production of sporopollenin, which forms part of the protective barrier for pollen (57). More work is needed to determine if the expanded family of *FAR2-like* genes identified here influences fertility or epicuticular wax levels in Chardonnay.

The Chardonnay genome enables thorough characterization of inter-clonal genetic variation. Attempts have been made in the past to use whole genome shotgun sequencing (WGS) to characterize inter-clonal diversity in other grapevine cultivars. These were ultimately limited by either available sequencing technology (58) or a lack of a reference genome for the particular grapevine variety under investigation (58, 59), although both studies were able to identify a small number of inter-clonal nucleotide variants. By taking advantage of both a reference genome for Chardonnay and increased read coverage, this study was able to identify 1620 high quality inter-clone nucleotide variants. There were limited shared somatic mutations among the Chardonnay clones, especially outside of the highly-related I10V1 group (I10V1, CR-Red and Waite Star). Clones 118 and 124, varieties from Burgundy used predominantly for sparkling wine production, were the exceptions to this, with 56% of their mutations being common between the two clones. Otherwise, the Chardonnay clones do not share a significant number of common mutations. This is likely the result of the centuries-long history of mass selection propagation. The clonal varieties of today likely represent a very small fraction of the genetic diversity that existed for Chardonnay after generations of serial propagation. The end result of this is that the many clones that were isolated from mass-selected vineyards appear to be genetically quite distinct from one another.

Furthermore, as the marker discovery pipeline developed in this study was limited in scope to detecting nucleotide polymorphisms within non-repetitive areas of the genome, there are likely to be structural variants, such as transposon insertions, that also impact on clonal-specific phenotypes. Nevertheless, marker mutations were identified for most of the clones that are predicted to impact gene function and could account for some of the clone specific phenotypes in Chardonnay.

Inter-clonal genetic variation provides an avenue for testing clone authenticity. The clone detection pipeline provides a fast and simple method to detect defined markers from a range of WGS library chemistries and platforms. Markers were reliably detected at coverages as low as 9.8-fold. Validation using independently-sourced clonal material indicated that most of the genetic variants were likely suitable for use in the identification of clones. Furthermore, there were a significant portion of markers that appeared to be variable across independently-sourced clonal material. This suggests that there might be region-specific genetic variation between clonal populations and this could potentially be exploited to further pinpoint the source of Chardonnay clones to specific regions, or to split clones into divergent subsets. The marker discovery and marker detection pipelines together form a solid framework for the future use of SNP- and InDel-based markers for the identification of unknown vegetatively propagated plant clones.

While the diploid Chardonnay reference genome enabled a much deeper understanding of the variation that has occurred since the initial establishment of this variety, it has also provided the means to unravel the detailed genetic ancestry of this variety and its parents, Pinot noir and Gouais blanc. Chardonnay matches both haplotypes of Pinot noir across approximately one fifth of its genome and these areas include large tracks of both homozygous and heterozygous variation. While the presence of the homozygous ‘double-Pinot noir’ regions could be result of a high number of large-scale gene conversion events early in Chardonnay’s history, the numerous heterozygous double-Pinot noir regions are only possible if the haplotype inherited from Gouais blanc was almost identical to the non-inherited allele of Pinot noir. Gouais blanc sequencing indeed confirms that within these ‘double-Pinot noir’ regions, one of the two Pinot noir haplotypes is a match for an allele of Gouais blanc.

The data reported in this work therefore supports a more complicated pedigree for Chardonnay than simply a sexual cross between two distantly related parents (Fig 6). The two parents of Chardonnay are predicted to share a large proportion of their genomes; this is suggestive of a previous cross between Pinot noir and a very recent ancestor of Gouais blanc (Pinot noir might even be a direct parent of Gouias blanc). Surprisingly, data supporting this complicated relationship between Gouais blanc and Pinot noir have appeared in previous low-resolution DNA marker analyses, with the two varieties sharing marker alleles at over 60% of marker loci in two separate studies (1, 31). However, the potential kinship between the two ancient varieties could not have been discovered without the insights provided by this diploid-phased Chardonnay genome.

**Fig 6:**
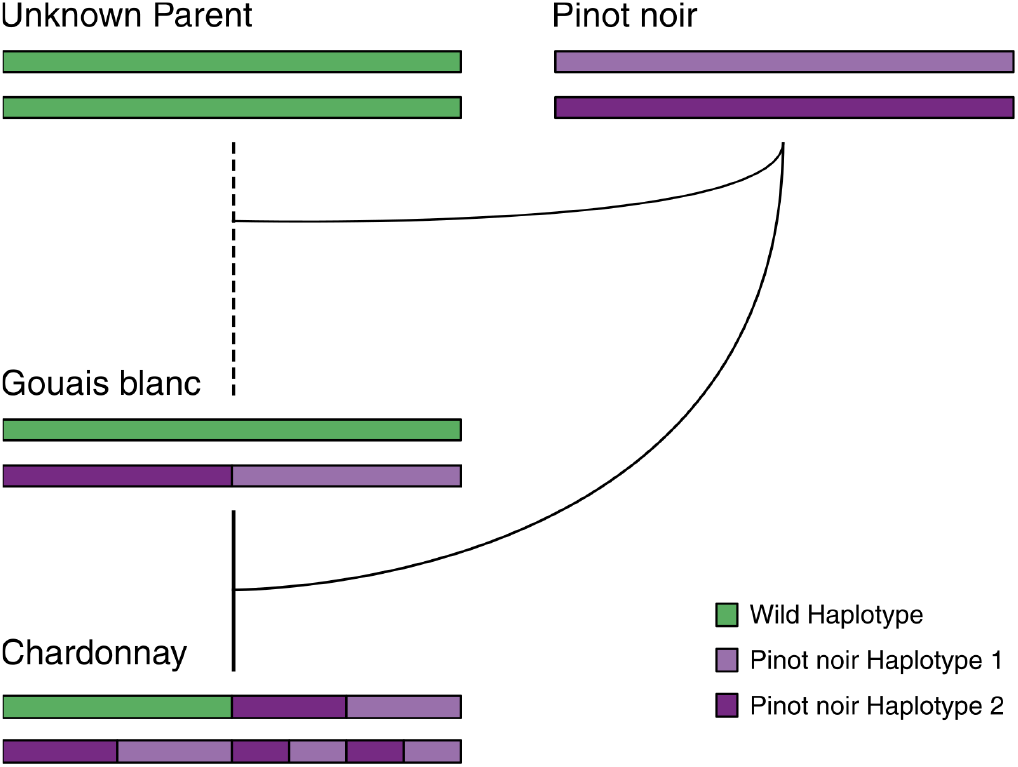
A schematic model for the complex pedigree of Chardonnay, Gouais blanc and Pinot noir. Two crossing events (akin to a standard genetic backcross) with Pinot noir would result in the homozygous and heterozygous Pinot noir regions present in Chardonnay.

A high-quality, diploid-phased Chardonnay assembly provided the means to assess several interesting facets of grapevine biology. It was possible to detect instances of heterosis, with differentially-expanded gene families being inherited from the parents of Chardonnay and to define the nucleotide variation that has accumulated during asexual propagation of this woody-plant species. However, most surprisingly, the completed genome indicates that the parents of Chardonnay shared a high degree of kinship, suggesting that the pedigree of this important wine-grape variety might be more complicated than originally thought.

## Methods

All custom scripts used for analysis, along with detailed workflows are available in S6 Archive. All sequencing data and the genome assembly have been lodged at the National Center for Biotechnology Information under the BioProject accession: PRJNA399599.

### DNA preparation and sequencing

Nuclear DNA was isolated from early season, disease free, field grown Chardonnay leaves taken from plants at a nursery vineyard (Oxford Landing, Waikerie, South Australia). DNA was extracted by Bio S&T (Quebec, Canada) from nuclear-enriched material using a CTAB/Chloroform method. DNA from clone I10V1 was enriched using a 1:0.45 Ampure cleanup prior to being used to build 15-50 kb SMRT Bell libraries with Blue Pippin size selection following library preparation (Ramaciotti Centre for Genomics, UNSW, Sydney, Australia). These libraries were sequenced on a PacBio RSII using 54 SMRT cells to give a total sequencing yield of 51,921 Mb (115-fold coverage) with an N50 length of 14.4 kb. Short-read sequencing of clones for marker discovery was performed on Illumina HiSeq 2000 and HiSeq X-Ten platforms from TruSeq libraries (100 and 150 bp paired end read chemistries). Short-read sequencing of clones for marker validation was performed on Illumina HiSeq 2500 and MiSeq platforms from Nextera libraries made from material sourced from both Foundation Plant Services (University of California, Davis) and Mission Hill Family Estate, Quail’s Gate and Burrowing Owl wineries in British Columbia, Canada.

### Assembly

The FASTA subreads were used to assemble the genome using FALCON (commit: 103ca89). Length cut-offs of 18 000 bp and 9 000 bp were used for the subread error correction and error-corrected reads respectively. FALCON Unzip (commit: bfa5e6e) was used with default parameters to phase the assembly from the FASTA subreads and Quiver-polish from the raw sequencing data.

The Purge Haplotigs pipeline (commit: f63c180)(29) was developed to automate the identification and reassignment of syntenic contigs from highly heterozygous long-read based assemblies. The PacBio RS II subreads were mapped to the diploid assembly (primary contig and haplotigs) using BLASR (packaged with SMRT-Link v3.1.0.180439)(60) and sorted with SAMtools v1.3.1. As required by Purge Haplotigs, read-depth thresholds were chosen to capture both peaks (diploid and haploid coverage levels) from the bimodal read-depth histogram and a contig-by-contig breakdown of average read-depth was calculated. Purge Haplotigs takes the read-depth summary and uses sequence alignments to reassign contigs. Curated primary contigs were assigned to *V. vinifera* chromosomes by using the PN40024 Pinot noir reference genome for scaffolding and for the identification of possible mis-assemblies. Several mis-assemblies were identified and manually corrected. The haploid and diploid curated assemblies were evaluated with BUSCO v3.0.1 using the embryophyta ODB v9 database.

### Annotation

A custom repeat library was produced for Chardonnay for use with RepeatMasker, similar to the method described in Fallon, Lower (61). Miniature inverted-repeat transposable element (MITE) sequences for *V. vinifera* were obtained from the P-MITE database (62). Repeats were predicted using RepeatModeler open-1.0.10 (63), and the RepeatModeler predictions and MITE sequences were concatenated to produce the custom Chardonnay repeat library. Repeats were annotated using RepeatMasker open-4.0.7 using this custom library.

RNA-seq was performed on total RNA extracted from I10V1 leaf tissue, extracted using a Spectrum Plant Total RNA Kit (Sigma), and sequenced using Illumina paired-end 75 bp chemistry on the Hiseq 2500 platform (Michael Smith Genome Sciences Centre, British Columbia Cancer Research Centre, British Columbia). Additional RNA-seq data from Chardonnay berry skins were obtained from the Sequence Read Archive (BioProject: PRJNA260535). All RNA-seq reads were mapped to the Chardonnay genome using STAR v2.5.2b (64), with transcripts predicted using Cufflinks v2.2.1 (65). Initial transcript predictions and repeat annotations were then used in the Maker gene prediction pipeline (v2.31.9) using Augustus v3.2.3 (66). The predicted proteins were assigned OrthoMCL (67) and KEGG annotations (68) for orthology and pathway prediction. Draft names for the predicted proteins were obtained from protein BLAST v2.2.31+ (69) hits against the Uniprot knowledgebase (70, 71) using an evalue cutoff of 1e-10.

### Parental mapping

Using BLAST and MUMmer v4.0.0beta (72) alignments, the primary contigs were aligned to the PN40024 reference, and the haplotigs were aligned to the primary contigs. The alignment coordinates were used to trim and extract the closely aligning phase-blocks between the primary contigs and haplotigs. BED files were produced that could be used for mapping the phase-blocks to the primary contigs and the chromosome-ordered scaffolds.

To identify the most likely parent for each phase-block pair, publicly-available short-read sequencing data were obtained for three clonally-derived variants of Pinot noir; Pinot blanc, Pinot gris, and Pinot meunier (BioProject: PRJNA321480); data for Pinot noir were not available at the time of analysis. To avoid potential issues with data from any single Pinot variety, pooled reads from all three were used for mapping. The sequencing data for Pinot, Chardonnay, and Gouais blanc were mapped to the primary contig and haplotig phase-block sequences using BWA-MEM v0.7.12 (73). PCR duplicates and discordantly-mapped reads were removed, and poorly mapping regions were masked using a window coverage approach. Heterozygous SNPs were called using VarScanv2.3 (p-value < 1e-6, coverage > 10, alt reads > 30%)(74) and Identity By State (IBS) was assessed over 10 kb windows (5 kb steps) at every position where a heterozygous Chardonnay SNP was found.

Where the parent (Pinot or Gouais blanc) was homozygous and matched the reference base, an IBS of 2 was called. Where the parent was homozygous and did *not* match the reference base, an IBS of 0 was called. Finally, where the parent and Chardonnay had identical heterozygous genotype, an IBS of 1 was called. The spread of IBS calls was used to assign windows as ‘Pinot’, ‘Gouais blanc’, or ‘double-match’. The window coordinates were transformed to chromosome-ordered scaffold coordinates and neighbouring identically called windows were chained together. Complementary Pinot/Gouais blanc calls from the parent datasets were merged and clashing calls removed. For ease of visualisation, the ‘double-match’ calls from the Pinot dataset were merged with the Gouais blanc calls (and vice versa). A SNP density track for the Chardonnay primary contigs was created over 5-kb windows from previously-mapped Illumina reads. The chromosome ideograms with SNP densities and IBS assignments were produced in Rstudio using ggplot2.

An orthologous kmer method for assigning parentage was developed to assign parentage over the entire genome. All 27-bp-long kmers (27mers) were counted using JELLYFISH v2.2.6 (canonical representation, singletons ignored)(75) directly from Pinot, Gouais blanc, and Chardonnay I10V1 PE reads to create 27mer count databases. Non-overlapping 1 -kb windows were generated for the primary contigs and the haplotigs. For each window all 27mers were extracted from the contig sequences, queried against the kmer count databases using JELLYFISH and the number of kmers not appearing in each were returned. The Pinot/Gouais blanc missing kmer counts were normalised against the Chardonnay counts and averaged over 10 kb windows with 5-kb steps. Windows with 150 or more missing kmers were classified as mismatch (missing kmer density was visualized over the genome to determine an appropriate cut-off) and neighbouring complementary windows were merged.

### Gene expansion

Protein-based BLAST alignments of Chardonnay proteins were performed against the Pinot noir (PN40024) reference proteome. Chardonnay Maker GFF annotations and PN40024 GFF annotations were converted to BED format. Chardonnay annotations were then transformed to scaffold coordinates. The ‘blast_to_raw.py’ script (from github.com/tanghaibao/quota-alignment) was used to flag tandem repeat homologues for both the primary contig and haplotig Chardonnay proteins against the Pinot noir reference. Illumina paired-end reads for Pinot were mapped to the Chardonnay primary and haplotig assemblies, BED annotations were created for regions with poor mapping, and these annotations were transformed to the scaffold coordinates for use with filtering. Predicted expanded gene families in Chardonnay that resided in Gouais blanc regions (identified using the kmer parental mapping) that had multiple gene models with poor read-coverage of Pinot mapped reads were returned as a filtered list. Dotplots were produced with MUMmer and the chromosomes were visually assessed for evidence of tandem sequence duplication at the filtered gene expansion candidate loci. The genomic sequences of *FAR2-like* ORFs from the Pinot noir assembly and from the Chardonnay primary contigs and haplotigs were aligned using MUSCLE v3.8.31 (76) within AliView v1.20 (77). Phylogenies were calculated within Rstudio using Ape (78) and Phangorn (79).

### Marker variant discovery

Paired-end reads for each clone were manually quality trimmed using Trimmomatic v0.36 (SLIDINGWINDOW:5:20, TRAILING:20, CROP:100)(80), mapped to the Chardonnay reference genome using BWA-MEM and filtered for concordant and non-duplicated reads. Variant calls were made using VarScan (p-value < 1e-3, alt reads > 15%) with variants across each clone pooled into a combined set. The combined variant set was then compared against leniently-scored variant calls for each clone (alt reads > 5, alt reads > 5%), with differences in genotype between clones resulting in that variant being flagged as a potential clonal marker.

Kmers were used to filter false positives from the pool of potential clonal markers. Kmer count databases (27mers) were created for each clone from the sequencing reads using JELLYFISH. For each potential marker, all possible kmers at the marker loci, from all samples, were extracted from the sequencing reads in the BAM alignment files. The kmer counts were queried from the kmer databases for each sample. Where a set of unique kmers were present for the matching samples, that variant was confirmed as a marker. The marker variants, marker kmers, and shared kmers were output in a table for use with querying unknown Chardonnay clones.

### Marker detection pipeline

Markers are detected directly from short-read sequencing data using kmers. A kmer count database (27mers) is calculated from raw sequencing reads as previously-described. The marker kmers and shared kmers that were identified in the marker discovery pipeline are then queried from the kmer database. A marker is flagged as a ‘hit’ if >80% of the marker kmers are present (at a depth of at least 3), and as ‘low-confidence hit’ if 30–80% of the marker kmers are present. Markers are flagged as ‘insufficient read coverage’ if fewer than 80% of the shared kmers are present at a depth of at least 12.

## Data Availability Statement

Raw sequencing reads, and the Chardonnay assembly and annotations are available under the BioProject accession PRJNA399599 at the National Center for Biotechnology Information. All custom code and analysis workflows required to reproduce the results presented are included in S6 Archive. Intermediate files can be made available on request.

## Acknowledgements

We thank Nick Dry (Yalumba Nursery) and Michael McCarthy (South Australian Research and Development Institute) for provision of plant material; Andrew Lonie and Torsten Seemann (University of Melbourne), Cihan Altinay and Derek Benson (University of Queensland), Stephen Crawley (QCIF), QRIScloud, and Melbourne Bioinformatics for assistance with computing resources; Jason Chin, Gregory Concepcion and Emily Hatas (Pacific Biosciences) for early access and guidance on FALCON Unzip; Sean Myles who acted as an advisor to Genome BC and Samantha Turner for outstanding administrative duties.

This project is supported by Australia’s grapegrowers and winemakers through their investment body Wine Australia, with matching funds from the Australian Government. Grapevine genomics work at the AWRI is also supported by Bioplatforms Australia as part of the National Collaborative Research Infrastructure Strategy, an initiative of the Australian Government. Funding was also obtained from Genome British Columbia and the Wine Research Centre at The University of British Columbia.

## Authors’ contributions

SAS, ARB, DLJ, HJJVV, SJJ, JB and ISP conceived and outlined the original approaches for the project. SAS sourced clonal material for reference sequencing and marker discovery. HJJVV, SJJ and JB provided material and sequence data for clonal validation. MJR, SAS and ARB designed and implemented the approaches used for genome assembly, marker discovery, and genome analysis. MJR, SAS and ARB wrote the manuscript, which was reviewed by all authors.

## Supporting information

**S1_Table.pdf: BUSCO analysis of the Chardonnay FALCON Unzip assembly before and after curation.**

**S2_Fig.pdf: Redundant primary contig reduction**. Circular representations of **A)** FALCON Unzip Chardonnay assembly and **B)** the same assembly after curation. Tracks are: length-ordered contigs **(i)**, read-depth of mapped Illumina paired end reads **(ii)** and heterozygous SNPs density **(iii)**.

**S3_Fig.pdf: Gene expansion of Chardonnay Chromosome 5 (primary contigs) region containing FAR2-like genes.** Alignments are indicated as black lines (dotplot), the ORFs for FAR2-like genes and pseudogenes are indicated for both Pinot noir and Chardonnay.

**S4_Fig.pdf: Distribution of clonal markers over the Chardonnay assembly**. Chardonnay chromosome-ordered primary contigs **(i)** and clonal marker variants **(ii)**.

**S5_Dataset.xlsx: Chardonnay clonal-specific markers. Sheet 1)** Chardonnay clone-specific markers with read-counts, **Sheet 2)** markers in gene models with Annovar and Provean predictions, **Sheet 3)** Summaries of clonal marker detection screening against validation datasets.

**S6_Archive.tar.gz: Scripts and workflows for data analysis.** Extract with tar for linux or Mac, or 7zip (7zip.org) for Windows. Contents: **bin/**, All custom scripts used for analysis; **lib/**, Custom Perl library for scripts; **src/**, Source code for window coverage masking program; **workflows/**, Commands used with comments for all data analysis; **Makefile**, The GNU Make pipeline for marker discovery.

